# N1 neuraminidase monoclonal antibodies 2H08 and 3H03 have decreased binding and inhibition properties against H1N1 influenza viruses circulating after 2018

**DOI:** 10.64898/2026.10.26.761829

**Authors:** Disha Bhavsar, Enikő Hermann, Brian Monahan, Jessica Nardulli, Charles Gleason, Viviana Simon, Florian Krammer

## Abstract

Influenza viruses are global public health problem. Vaccines are available but have to be updated annually due to antigenic drift of the main vaccine component, the surface glycoprotein hemagglutinin (HA). The second surface glycoprotein of the virus, neuraminidase (NA), has recently moved into the spotlight of vaccine development. However, its antigenic evolution is understudied. We found that two broadly protective N1 NA antibodies, which had previously been shown to bind and inhibit most N1 NA, lost activity to recent N1s from seasonal H1N1 viruses. Here, we show that this drop in activity can mostly be attributed to a change in electrostatic surface potential from neutral to positively charged. This effect was not observed when historic human plasma samples were tested, suggesting that the drop of activity is specific to the two tested mAbs but does not broadly impact the polyclonal human response. Finally, we confirmed that the tested mAbs were fully active against recently emerged panzootic clade 2.3.4.4b H5N1 NA.

**Importance:** The influenza virus NA has become an important target for vaccine development. The characterization of antigenic changes described in this manuscript contributes to our understanding of conservation of epitopes in NA.

## Introduction

Seasonal influenza virus infections cause significant morbidity and mortality every year. Vaccines against these viruses are mostly based on driving an immune response against the virus hemagglutinin (HA). HA mediates binding of the virus to cells via terminal sialic acid on N-linked glycans and fusion of the viral membrane and the cellular endosomal membrane to promote release of the viral genome into the cytoplasm (1). Due to these important functions, antibodies to HA are highly neutralizing. However, influenza viruses evolve at a high rate and due to antigenic drift of HA, available vaccines have to be updated and readministered annually (2). The second surface glycoprotein, the neuraminidase (NA), is an enzyme that cleaves terminal sialic acid from N-linked glycans (3, 4). This receptor destroying activity is important during early stages of infection when the virus needs to pass through mucosal fluids to reach host cells. Mucosal fluids contain highly sialilated natural defense proteins like mucins which can trap virus. NA activity frees the virus from these interactions (5). NA is also very important in a much later stage of virus replication, as it allows virus to be shed from infected cells (6). Without NA activity, newly released virus would get stuck on sialic acid on infected cells and would not be able to disseminate. NA also undergoes evolution and antigenic drift (7-10), but much less is known about this process compared to HA antigenic drift. However, conserved epitopes are present on NA and NA has become a focus for development of broadly protective vaccines in recent years (11-16).

We have isolated two broadly protective anti-N1 monoclonal antibodies (mAbs)from a survivor of the 2009 H1N1 pandemic (13). Our initial findings showed, that these antibodies bound to most tested N1 NAs including N1s from seasonal, zoonotic and animal viruses (13). However, we have noticed that the activity of these mAbs is diminished or absent for more recent seasonal H1N1 strains. Here, we investigate the molecular determinants of this loss of activity.

## Results

### Broadly inhibiting anti-N1 mAbs show decreased binding and inhibition against recent influenza virus N1s

Recently, Hansen and colleagues (17) identified and characterized two broadly inhibiting mAbs, 2H08 and 3H03, against the N1 NA, isolated from a patient infected with the 2009 pandemic H1N1 influenza virus. Their experiments have shown that the antibodies were able to successfully bind to a range of N1 NAs and inhibit the activity of the NA. The mAbs were also able to prevent fatal outcomes of infection *in vivo* against both H1N1 isolates of human origin, as well as H5N1 avian influenza virus strains. This study tested HXN1 strains from the years 1918-2015.

However, preliminary measurements with H1N1 strains isolated after 2020 indicated a decrease in binding and inhibition efficiency of these mAbs. Therefore, we set out to measure the properties of mAbs 2H08 and 3H03 *in vitro* against various H1N1 or H5N1 isolates from 2009-2024 (**Figure 1**), focusing mainly on H1N1 influenza virus vaccine strains as representative viruses. Our measurements showed that there is indeed a decrease in binding activity measured by enzyme-linked immunosorbent assays (ELISA), and also inhibition activity, as measured by enzyme-linked lectin assays (ELLA), for isolates from 2018-2022. This was especially true for 3H03, but 2H08 also showed decreased inhibition activity against certain isolates sampled after 2019. Notably, for historical and recent H5N1 isolates from clade 2.3.4.4b strong binding in ELISA measurements was maintained.

**Figure 1.**
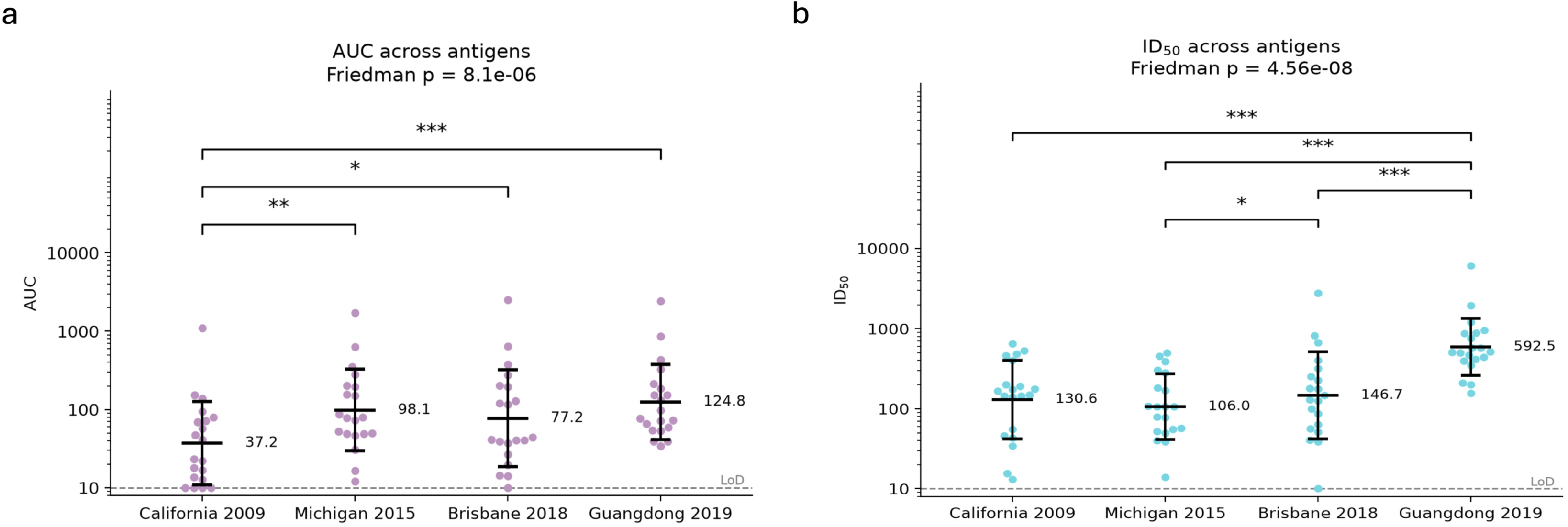
Combined phylogenic tree and heat map representing the different influenza virus isolates used for analysis and binding and inhibiting activity of the tested mAbs. The combined phylogenic tree and heat map was prepared with *in vitro* binding activities of mAbs 3H03 and 2H08 measured using ELISA (blue) and *in vitro* inhibition activity of mAbs 2H08 and 3H03 as measured by ELLA (purple) against N1 antigens. NAI measurements marked with a star had detectable inhibition in ELLA measurements, however, IC_50_ was >30 μg/mL.

To summarize, two broadly inhibiting anti-N1 mAbs, 2H08 and 3H03, showed decreased binding and inhibition efficiency towards seasonal H1N1 influenza viruses isolated from humans after 2018.

### Accumulating mutations at the epitope of 2H08 and 3H03 lead to a change of electrostatic surface potentials

The exact epitopes of mAbs 2H08 and 3H03 were known, as experimental cryo-electron microscopy structures were available (PDB codes: 8E6K and 8E6J respectively). This gave us the opportunity to pinpoint the sites that mutated between 2009 and 2024 within the epitope of the mAbs. We prepared models for all the representative N1 head domains using Chai-1 (18) to be able to identify changes. We could identify four key mutations occurring sequentially at the epitopes of 2H08 and 3H03 in this period: N386K, N449D, D416N and I389K (**Figure 2a and c)**. These mutations occur for the first time in the representative strains in the Northern hemisphere influenza seasons 2015-2016, 2017-2018, 2018-2019 and the Southern hemisphere season 2019 respectively.

**Figure 2.**
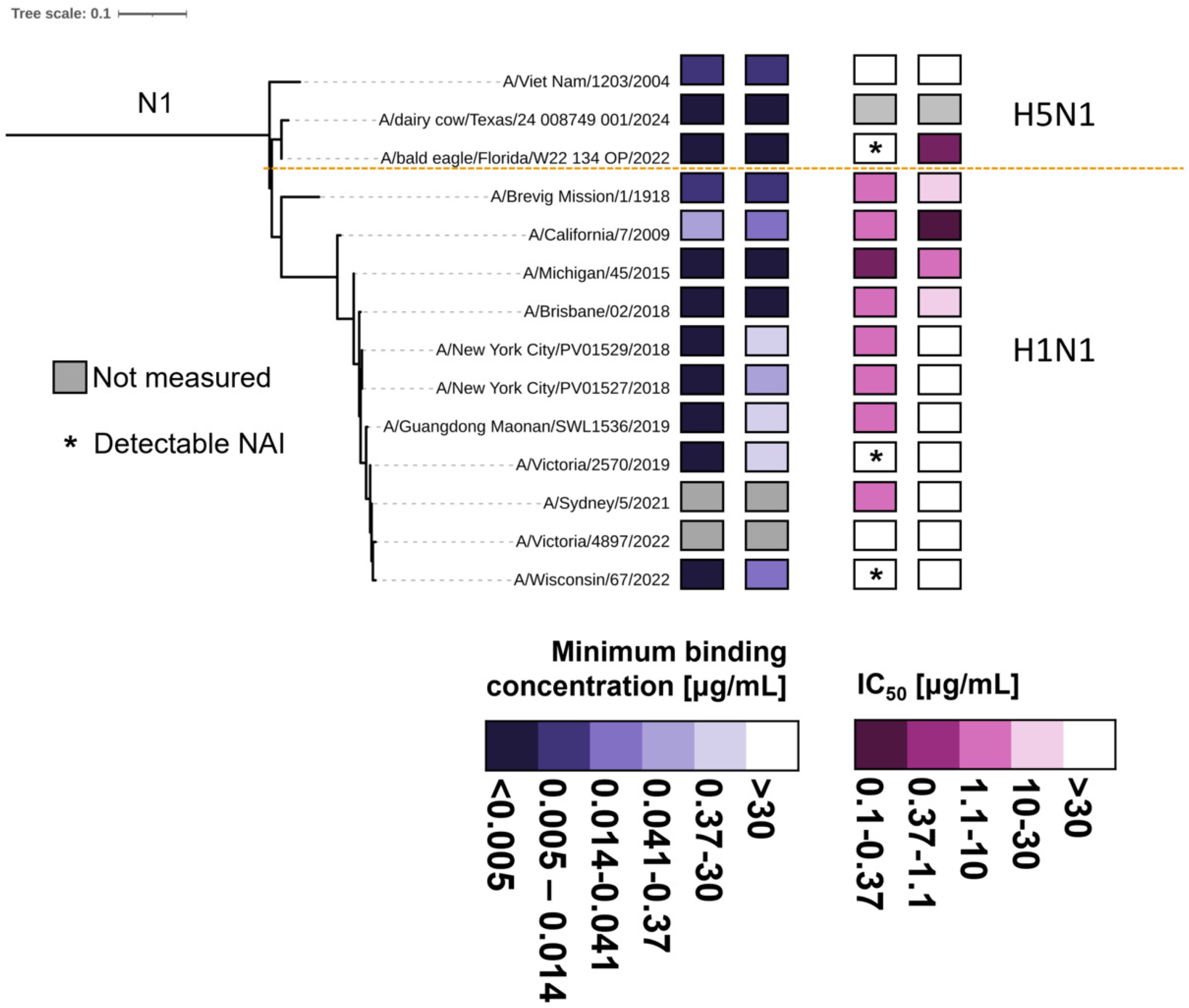
Accumulating mutations in the epitope of mAbs 2H08 and 3H03 cause a change in electrostatic surface potential. **A**) Aligned models of the N1 neuraminidases from the isolates A/California/04/2009 (orange) and A/Victoria/2570/2019 with the mAb 2H08 (light pink cartoon). The key mutations occurring on the epitope site are shown as sticks. **B**) Electrostatic surface potentials of mAbs 2H08 and 3H03. The yellow outline shows the paratope areas. **C**) Change of electrostatic surface potentials in the epitope of 2H08 and 3H03 in structural models of NAs of representative vaccine strains between 2009-2022. The location of the epitope is marked orange on the VASP tetramerization domain – NA head model of A/California/04/2009.

The main change these mutations cause is in the electrostatic surface potential of the epitope (**Figure 2c**). The paratopes of both mAbs are mostly positively charged (**Figure 2b**), while the epitope of A/California/04/2009, the virus the mAbs were likely first induced against, is negatively charged (**Figure 2c**). The same is true for the N1 epitopes of the H5N1 representatives measured in wet-laboratory experiments (**Figure S1**). Two of the four mutations accumulating on the epitope site change neutral amino acids to positively charged ones (N386K and I389K), and one changes a negatively charged one to neutral (D416N). The exception is N449D, that adds a negatively charged aspartate to the epitope surface. Of note, the recommended vaccine strain for the 2026/2027 Northern hemisphere season, A/Missouri/11/2025 (H1N1), has the same four mutations at the epitope comparted to A/California/04/2009.

In conclusion, the mutations occurring sequentially at the epitope surface change the electrostatic surface potential of the epitope from negative to positive, which makes it incompatible to bind the positively charged paratopes of the mAbs. N1s from historical and recent H5N1 avian influenza virus strains have a negatively charged epitope, allowing binding.

### Accumulated mutations influence molecular interactions between mAbs 2H08 and 3H03 and N1s from human influenza viruses

Next, we aimed to investigate in detail the contributions of the mutated amino acids on N1 to the antibody-antigen interactions.

For this, we used molecular dynamics simulations. In these simulations, we included NA of the A/California/04/2009 and A/Victoria/2570/2019 H1N1 influenza viruses and used a vasodilator-stimulated phosphoprotein (VASP)-tetramerization domain (19) instead of the stalk domain, to match the experimental ELISA measurements. The simulations also included Ca^2+^-ions and glycosylations at the predicted N-glycosylation sites. Both viral NAs were simulated as tetramers, and each tetramer had a mAb attached. Separate simulations were done for all combinations of NAs and mAbs, giving a total of 4 NA-head mAb complex to evaluate. This meant a total of 4x100 ns (0.4 us) simulation for each system.

The first mutation, occurring from 2015 onward is N386K, an asparagine mutating to a lysine. This mutation changes the site from neutral to positively charged. In addition, it also removes a glycosylation site near the mAbs’ epitope (**Figure 3a**). On a molecular level, this influences the hydrogen bonding capabilities of this residue. In one of the tetramers, N386 makes a hydrogen bond with T32 of the mAb 2H08 in 44% of the simulation snapshots, and this interaction is also present sporadically in the other monomers. With the mutation to K386, this interaction completely disappears from any of the monomers (**Figure S2**). No such molecular effect was observed towards 3H03.

**Figure 3.**
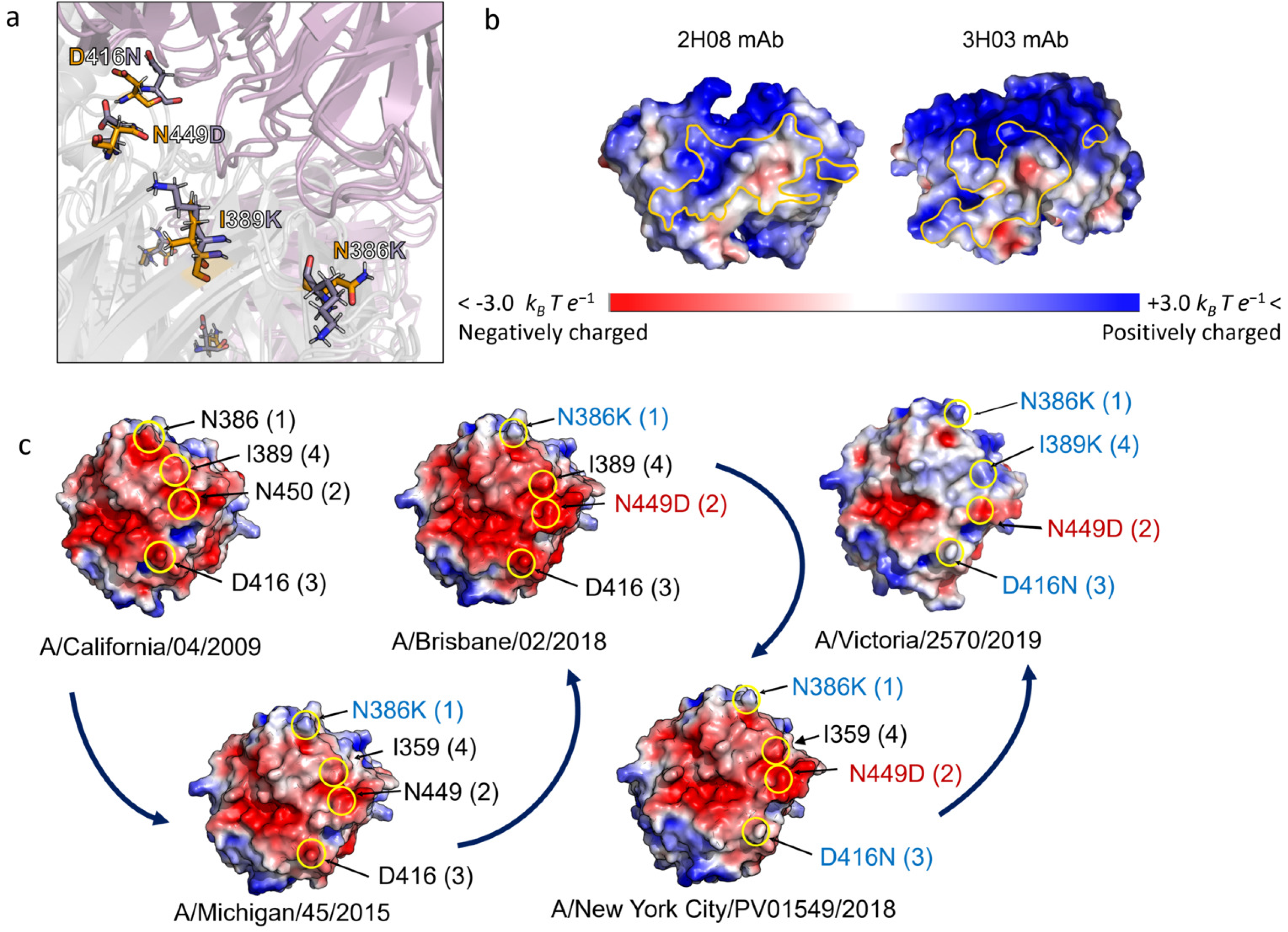
Key interactions between mAbs 3H03 and 2H08 and A/California/04/2009 (orange) or A/Victoria/2570/2019 (purple) NA. **A**) Hydrogen bonding interactions made by N/K386 with T32 of the heavy chain of 2H08. **B**) Hydrogen bonding interactions made by N/D449 with T53, S52 and Y49 of the heavy chain of 2H08. **C**) Hydrogen bonding interactions made by D/N416 with S106 of the heavy chain of 3H03. **D**) Salt bridge interactions made by D451 with R56 of the light chain of 2H08.

The second mutation appearing in 2018 is N449D. This is the only mutation that adds a negative charge instead of a neutral. N449 makes key hydrogen bonding interactions with Y49 and T53 of 2H08 and I104 and S105 of 3H03 (**Figure 3b**). The mutation to D449 does slightly influence these hydrogen bonds: they remain present majorly towards T53 and S52 for 2H08 and T102 and S105 for 3H03. However, the total number of hydrogen bonds throughout the simulation are even higher with D449 than with N449 for both mAbs (**Figure S3**).

The third mutation appearing in 2018 is D416N. This changes a negatively charged residue to a neutrally charged one. It influences the binding of 3H03 to the NA heads: While D416 readily makes 1-2 hydrogen bonds in all snapshots with S106 and N108 in 3H03, these bonds are present more rarely throughout the simulation with N416 (**Figure 3c**). D416 is contributing to at least one hydrogen bond throughout 73-90% of the simulation time; while for N416 these ratios are 1-29%, depending on the monomer. Also, N416 is able to make one hydrogen bond, at one timepoint, while D416 often contributes to two (**Figure S4**).

The I389K mutation is again one that changes a neutrally charged amino acid to a positively charged one. This residue does not contribute to hydrogen bonding interactions, but likely is a major contributor to the overall charged-charged repulsions described in the previous section.

The main salt-bridge forming residues on A/California/04/2009 NA are D384, forming a bridge with R56 in the heavy chain Fc of 2H08, and D451, with R56 in the light chain of 2H08 (**Figure S5**). While these residues did not mutate between 2009-2022, they are neighboring residues to N386K and N449D. Notably, the D384-R56 heavy chain interactions are near the glycosylation on N386, which most likely sterically prevents a hydrogen bonding interaction. This interaction changes to a hydrogen bonding interaction for D384 in A/Victoria/2570/2019 (**Figure S6**). D384 also is a coordinating residue for a Ca^2+^-ion. D451 forms a salt-bridge with R56 in the light chain. This salt bridge is present in more of the snapshots overall for A/California/04/2009 than for A/Victoria/2570/2019 (**Figure S5, Figure 3d**). However, no salt bridges or charged-charged repulsive interactions were identified for N1s from A/California/04/2009 or A/Victoria/2570/2019 when simulated together with 3H03.

To summarize, the four mutated residues at and near the epitope sites influence not only the electrostatic surface potentials, but also hydrogen bonding and salt-bridge interactions between the N1s and mAbs.

### “State-of-the-art” methods for predicting protein-protein structures fail to predict correct orientation of NA antigen – antibody complexes

In the A/California/04/2009 and A/Victoria/2570/2019 NA – 2H08, 3H03 mAb antigen complex systems we have knowledge of the exact epitope: an experimental structure has been solved for both of these mAbs together with influenza A H1N1 virus NA from A/Brevig Mission/1/1918 (PDB: 8E6J and 8E6K). This gives a good comparison point for any models, and also makes it possible to model other NAs together with 2H08 and 3H03 by aligning them to the solved structures.

Several advances have been made for predicting structures of protein-protein complexes in the last five years. Here, we attempted to create models of the antigen-antibody complexes of A/California/04/2009 and A/Victoria/2570/2019 N1s together with the mAbs 2H08 and 3H03 using ESMFold2, and Chai-1, Boltz-2, the latter two both with and without residue-residue restraints for correctly orienting the mAbs. We did not include AlphaFold3 as the strict licensing does not permit the use of outputs in docking or screening tools. As a reference, we chose the models created by aligning the different NA models to our reference experimental structures 8E6J and 8E6K, and including the antibodies in this orientation together with the NA models (see Materials and Methods for details). Then we calculated DockQ of the NA head – antibody heavy chain scores to find the best modeling method (**Table 1**).

**Table 1.** DockQ scores of the NA– mAb structures compared to models based on the experimental structures. DockQ scores are interpreted as: < 0.23 – Incorrect (red); 0.23 ≤ Acceptable < 0.49 (orange); 0.49 ≤ Medium quality <0.80 (light green); ≥ 80 High quality (dark green).

|  | Boltz-2 no restraints | Boltz 2 with restraints | Chai-1 no restraints | Chai-1 with contacts restraints | Chai-1 with pocket restraints | ESMFold 2 |
| --- | --- | --- | --- | --- | --- | --- |
| A/California/04/2009 with 2H08 | 0.14 | 0.23 | 0.006 | 0.86 | 0.33 | 0.018 |
| A/California/04/2009 with 3H03 | 0.054 | 0.061 | 0.11 | 0.87 | 0.36 | 0.13 |
| A/Victoria/2570/2019 with 2H08 | 0.027 | 0.24 | 0.11 | 0.63 | 0.26 | 0.008 |
| A/Victoria/2570/2019 with 3H03 | 0.065 | 0.057 | 0.33 | 0.83 | 0.14 | 0.008 |

As visible, most “state of the art” protein-protein structure prediction methods still struggle to predict antigen-antibody interactions correctly. This is improved by adding distance restrains with Chai-1. However, this method needs input with exact residue-residue contacts which is hard to obtain without using experimental structural biology methods. The “pocket” restraints of Chai-1 are equivalent to using Boltz-2 with restraints: both only need to define residues on just the antigen where the antibody attaches, and define the attaching antibody chain. These methods are moderately successful in finding the correct positions of antibodies. This information is obtainable by creating escape mutants in wet-laboratory experiments, without the need of costly structural biology experiments (20, 21). However, as show, this information is not necessarily enough for creating reliable antibody-antigen structures with current artificial intelligence (AI) based structural biology methods.

### Human polyclonal sera from before 2018 retains binding and inhibition towards NAs from strains isolated after 2019

Finally, we aimed to investigate if these mutations influence the influenza virus NA binding and inhibition properties of human polyclonal antibodies. For this we performed ELISA and NAI experiments using 20 human polyclonal plasma from adult study participants vaccinated against influenza in the 2016-2017 season (**Figure 4**) prior to the emergence of viruses carrying mutations causing changes in the binding and inhibition of the mAbs 2H08 and 3H03. While we saw significant differences between the binding and inhibition of different influenza virus N1s by these biospecimen, there was no decrease of binding or inhibition with the N1s sampled after 2018. The inhibition efficiency even increased against A/Guangdong Maonan/SWL1536/2019, which features four mutations on the epitope of 2H08 and 3H03. These data indicate that despite the change of one epitope site, human polyclonal responses remain active against influenza virus N1 despite the changes in the 2H08/3H03 epitope.

**Figure 4.**
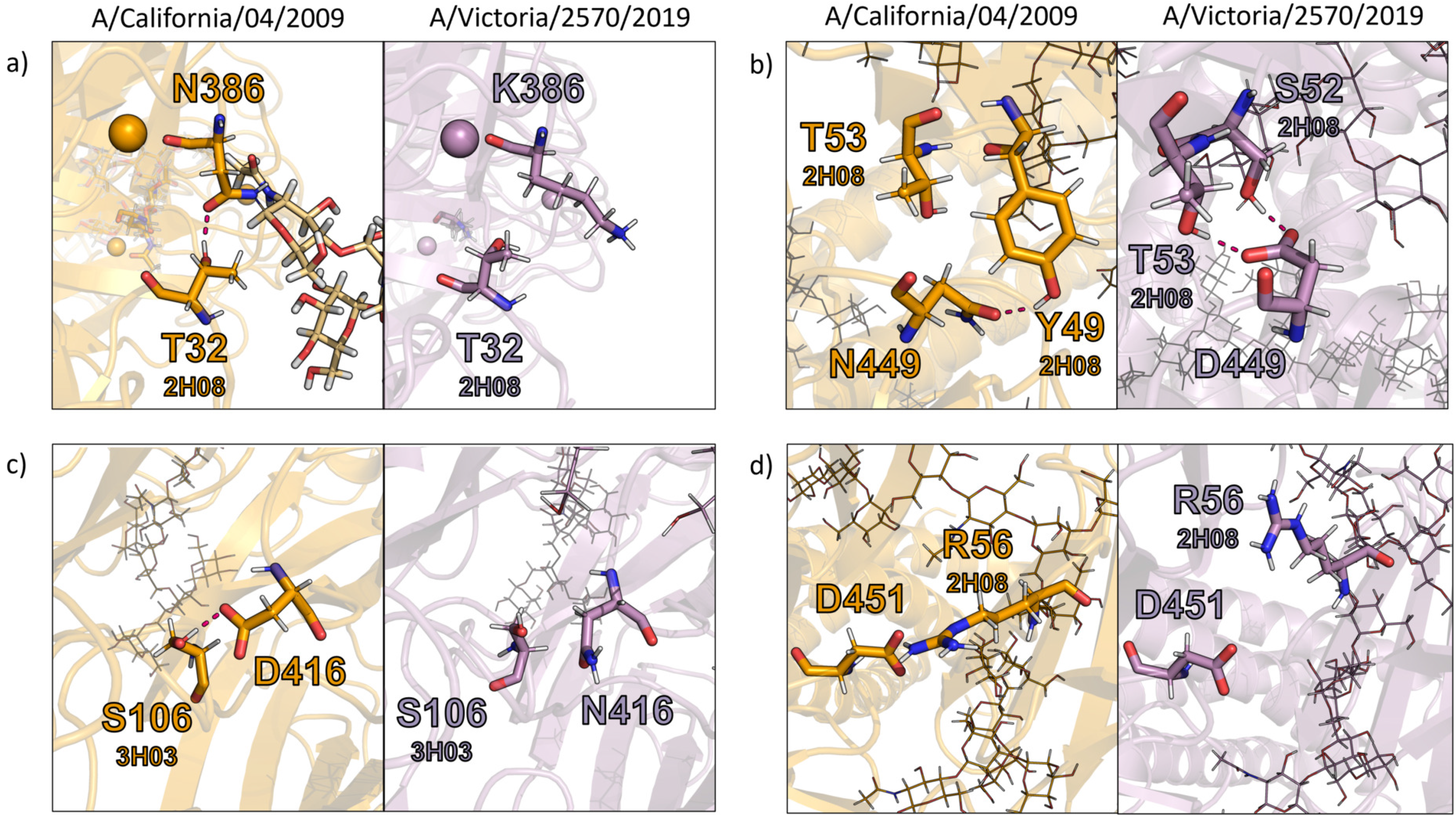
Binding and inhibition activity of human polyclonal plasma samples from 2017 against various human N1s. **A**) Immunoglobulin G (IgG) antibodies binding to recombinant N1 proteins from A/California/04/2009, A/Michigan/45/2015, A/Brisbane/02/2018 and A/Guangdong Maonan/SWL1536/2019 H1N1 viruses measured using ELISA. Binding titers are presented as area under the curve (40) with error bars representing geometric mean and standard deviation and the limit of detection is represented as a dotted line. **B**) Neuraminidase inhibiting antibody responses in plasma against the panel of H1N1 viruses were measured using ELLA. NAI titers are shown as ID_50_ with error bars representing. Geometric mean with standard deviation and limit of detection is represented as a dotted line. Pairwise Wilcoxon signed-rank tests (paired) between every antigen pair with Holm-Bonferroni correction was used for pairwise comparison of the plasma samples against different N1 antigens.

## Discussion

Antibody titers to NA have been recognized as independent correlate of protection and NA has also become a prime focus of influenza virus vaccine development (15, 22-26). A number of broadly protective antibodies to NA have been identified that target conserved epitopes (12-14, 16, 27-30). In addition, vaccine studies with NA immunogens have demonstrated that these immunogens can induce broad protection within a subtype (31-37). However, it is clear that NA also undergoes antigenic drift (7-10).

Here, we focused on understanding the conservation of the epitopes of two broadly protective anti N1 mAbs, 3H03 and 2H08. We have previously shown that these mAbs target a broad range of diverse N1 NAs (13). In preliminary studies, we saw a decrease in binding of these two antibodies to recent H1N1 isolates. Here, we investigated the molecular basis for this loss of activity using a detailed analysis of the epitope-paratope interaction and changes in the epitope over time. We found that mutations in the epitope caused the surface to become increasingly positively charged which resulted in weaker interactions between the antibodies and the N1 NA. Importantly, binding and neuraminidase inhibition of recent clade 2.3.4.4b H5N1 viruses was fully retained.

To investigate if these changes were specific to the mAbs or if they indicated broader antigenic drift, we also investigated binding and inhibiting activity of human plasma collected before these changes in the NA occurred. With polyclonal antibody no decrease in binding was observed. In fact, binding to more recent viruses was slightly higher and NI activity to more recent viruses slightly increased as well. This suggests that the observed changes do not represent *bona fide* antigenic drift.

We used modeling combined with molecular dynamics (MD) simulations to investigate the individual contributions of amino acid changes on the epitopes of the mAbs 2H08 and 3H03. With the appearance of the third mutation, D416N in 2018, we see a drop in the binding and inhibition capabilities of 3H03 but not 2H08. This is in line with our molecular level observations in MD simulations: with this mutation the hydrogen bonding interactions made by N416 decrease compared to D416, but only for mAb 3H03. For a decrease in inhibition of the mAb 2H08 an additional mutation, I389K is necessary, which changes a neutral, apolar isoleucine to a positively charged lysine. While this mutation does not influence the hydrogen bonding capabilities, it changes a large negatively charged patch on an epitope site to a positively charged one. This, together with the decreased hydrogen binding effects of N386K results in also decreased *in vitro* inhibition capabilities by 2H08. However, the first mutation itself, N386K was not sufficient for the same effect. Notably, no charged-charged interactions were identified involving any of the residues mutating to a changed charged one. It is likely that the change of the electrostatic surface potentials presents as an overall effect in binding, rather than influencing the interaction between two amino acids of the N1 and the mAbs.

Since the publishing of AlphaFold2 in 2021, exponential progress has been made in the field of AI-based structure prediction. New models become available regularly, improving on prediction quality, speed, and types of molecules that can be included. We tested the popular models ESMFold2, Chai-1 and Boltz-2. However, antibody-antigen complexes remain difficult to correctly predict. Our findings confirm this claim: without detailed wet-lab experiment-based input, none of the tested models managed to successfully predict the orientation of 2H08 and 3H03. This is reflected also in the prediction quality metrics: the ipTM scores between the NA and the mAb chains remain consistently below the acceptable 0.6. The only exception is the usage of Chai-1 with residue-residue restraints. However, this requires prior knowledge of the exact epitope site and available interactions. This is in line with the findings of benchmarking efforts for structures of antibody-antigen complexes: as of September 2026, even with the best available methods only 18.4% of the predictions is scored as a high-quality model on CoFold Arena (38). Therefore, we advise to use AI-based structure prediction methods with caution when predicting the possible interactions of new mAbs with their targets, and also careful observation of confidence metrics when reporting such findings.

In summary, we show that changes in surface charge led to the loss of activity of mAbs 3H03 and 2H08 against recent H1N1 strains. However, the changes in this conserved epitope do not seem to impact activity of polyclonal human plasma and therefore do not seem to indicate proper antigenic drift.

## Materials and Methods

### Study participant details

We selected biospecimen from 20 adult study participants of an observational longitudinal institutional review board (IRB)-approved cohort study conducted in New York City (STUDY-16-01199). All participants provided written informed consent prior to data/sample collection. Plasma samples were collected in 2016/2017 and stored at -80ºC until use.

### Cells, viruses and recombinant proteins

*Spodoptera frugiperda* (Sf9) cells were grown in *Trichoplusia ni* medium-formulation Hink (TNM-FH) insect cell medium (GeminiBio) supplemented with 10% fetal bovine serum (FBS; Gibco) and antibiotics (100 U/ml of penicillin, 100 µg/ml streptomycin; Gibco). High Five cells (BTI-TN-5B1-4; Boyce Thompson Institute-*Trichoplusia ni*-5B1-4 subclone; Vienna Institute of Biotechnology) cells were grown in serum free Sf-900-ii medium (Gibco) supplemented with antibiotics (100 U/ml of penicillin, 100 µg/ml streptomycin; Gibco). Expi293 cells (Gibco# A14527) were grown in Expi293 medium (Gibco) in suspension shaking at 125 RPM in a humidified incubator at 37°C with 8% CO_2_.

Recombinant NA proteins were expressed and purified using baculovirus-insect cell expression system as previously described in detail (39). Briefly, the ectodomains of the NA proteins were cloned into a baculovirus shuttle vector containing an N-terminal signal peptide followed by a hexahistidine tag, a vasodilator-stimulated phosphoprotein (VASP) tetramerization domain and a thrombin cleavage site. The shuttle vectors were used to generate baculoviruses which were propagated in Sf9 cells and used to infect High Five cells for protein expression. Seventy-two hours post infection, the cell culture supernatants were harvested, and the recombinant NA proteins were purified using affinity purification with nickel-nitrilotriacetic acid (Ni-NTA) agarose beads (Qiagen).

Influenza viruses were grown in 10-day old specific pathogen free (SPF) embryonated chicken eggs (AVSBio) for 48 hours at 37°C and cooled at 4°C overnight. The allantoic fluids were collected and clarified by centrifugation at 300x*g* for 5 minutes at 4°C. The virus stocks were aliquoted and stored at -80°C until further use.

Plasmids expressing heavy and light chains of the mAbs were transiently transfected into Expi293 cells using an Expi293 transfection kit per manufacturer’s instructions. The supernatants were collected at 7 days post transfection and the antibodies were purified using a CaptureSelect IgG-Fc Multispecies Affinity Matrix (Thermo Scientific).

### Plasma sample desalting

Prior to use in assays, human plasma were heat inactivated at 56°C for 1 hour and desalted using Zeba™ Spin Desalting Columns, 40K molecular weight cutoff (Thermo Scientific) per manufacturer’s instructions to remove ethylenediaminetetraacetic acid (EDTA).

### ELISA

96-well plates (Immulon-4 HBX; Thermo Scientific) were coated with 50 µl/well of recombinant NA protein at a concentration of 2 µg/ml and stored at 4°C overnight. The following day, the coating solution was discarded and the plates were blocked with 200 µl/well of 3% w/v non-fat milk prepared in phosphate buffered saline (PBS) with 0.1% Tween-20 (PBS-T) for 1 hour at room temperature (RT). MAbs were diluted at a starting concentration of 30 µg/ml and further diluted 1:3 serially in 1% non-fat milk prepared in PBS-T. For the ELISA performed with plasma, samples were diluted 1:50 initially followed by 1:2 serial dilutions. Following the incubation, the blocking solution was removed and 100 µl of diluted mAbs or plasma were added to the corresponding wells and incubated at 2 hours at RT. Broadly reactive neuraminidase antibody 1G01 (30) was used as a positive control and a Severe acute respiratory syndrome coronavirus-2 (SARS-CoV-2) mAb was used as a negative control. The plates were then washed 3 times with 200 µl/well of PBS-T and 50 µl of secondary antibody (Goat anti−human IgG (Fab-specific)−horseradish peroxidase (HRP) labeled antibody; Sigma) was added to each well. The plates were incubated for 1 hour at RT, and washed with 200 µl/well of PBS-T. Following this, 100 µl/well of SigmaFAST OPD (o-phenylenediamine dihydrochloride; Sigma-Aldrich) was added to the plates and incubated for 10 minutes. The reaction was stopped using 50 µl/well of 3M hydrochloric acid (HCl) and the optical density (OD) at 490nm was recorded using SynergyH1 microplate reader (BioTek). For mAbs, minimal binding concentration was determined as the lowest concentration at which the OD value was higher than baseline (defined as average plus 3-times standard deviation of blank wells). Area under the curve (40) values for plasma samples were calculated using GraphPad Prism 11 software.

### ELLA

ELLAs were performed to first assess the neuraminidase activity of the virus or recombinant NAs. 96-well plates (Immulon-4 HBX, Thermo Scientific) were coated with 100 µl/well of 25 µg/ml fetuin (Sigma) diluted in PBS and stored at 4°C overnight. The following day, the plates were washed 3 times with 200 µl/well of PBS-T. Two-fold serial dilutions of virus starting at 1:10 initial dilution or recombinant NA starting at 30 µg/ml prepared in PBS-T supplemented with 1% bovine serum albumin (BSA) were added to the plates and incubated overnight at 37°C. The plates were then washed 6 times with 200 µl/well of PBS-T and 100 µl of peanut agglutinin conjugated with horseradish peroxidase (PNA-HRP; Sigma) diluted to 5 µg/ml in PBS supplemented with 1% BSA was added to each well and incubated for 2 hours at RT. Plates were then washed 3 times with 200 µl/well of PBS-T. Hundred µl/ well of SigmaFAST OPD was added to the plates and incubated for 10 minutes at RT. Fifty µl/well of 3M HCl was used to stop the reaction and OD at 490nm was measured using SynergyH1 microplate reader. The data were analyzed using GraphPad Prism 11 and 50% effective concentration (EC_50_) was determined.

To measure the neuraminidase inhibition activity, either mAbs serially diluted 1:3 at a starting concentration of 30 µg/ml, or plasma serially diluted 1:2 after 1:10 initial dilution were added to fetuin coated plates. An equal volume of virus or recombinant NA diluted to 2xEC_50_ were added to the dilutions and incubated overnight at 37°C. The remainder of the assay was performed as described above. Broadly reactive neuraminidase antibody 1G01 (30) was used as a positive control and a SARS-CoV-2 mAb was used as a negative control. Percent inhibition of neuraminidase activity was determined and 50% inhibitory concentration (IC_50_) values for mAbs or 50% inhibitory dilution (ID_50_) values for plasma were calculated using GraphPad Prism 11.

### Statistical analysis

To analyze the overall differences in AUC and ID_50_/IC_50_ obtained from ELISA and ELLA measurements of human plasma samples, the Friedman test was used. Pairwise Wilcoxon signed-rank tests (paired) between every antigen pair with Holm-Bonferroni correction was used for pairwise comparison of the plasma samples against different N1 antigens. Results were considered significant if p<0.05. Python libraries scipy (41), pandas (42) and numpy (43) were used for calculations and libraries matplotlib (44) and seaborn (45) for plotting.

### Structural model building, structural analysis and molecular dynamics simulations

Initial models of the N1 heads of A/California/04/2009, A/Michigan/45/2015, A/Brisbane/02/2018, A/New York City/PV01549/2018 and A/Victoria/2570/2019 were predicted as tetramers using Chai-1 on the Chai-1 server (18). These models were used to calculate the electrostatic surface potentials of chain A of the models with the Adaptive Poisson-Boltzmann Solver (APBS) (46).

Chai-1 was also used to create models of A/California/04/2009 and A/Victoria/2570/2019 including the VASP tetramerization domains with NA heads, matching experimentally used constructs. The heads of the A/California/04/2009 or A/Victoria/2570/2019 N1 tetramers were first aligned using PyMol (47) with PDB: 8E6K or PDB: 8E6J, and the 2H08 and 3H03 mAbs were included from these alignments to get full models including the antibodies. GLYCAM-Web (48) was used to add M5 glycans (Man(a1-3)[Man(a1-3)[Man(a1-6)]Man(a1-6)]Man(b1-4)GlcNAc(b1-4)GlcNAc) at each predicted N-glycosylation site. The full, glycosylated models were aligned to the PDB: 3NSS structure to include a total of 9 Ca^2+^-ions. These models were then used as input into molecular dynamics (MD) simulations. Simulations were carried out using OpenMM (49), with the Amber14 ff14SB (50) and GYLCAM06 J-1 (51) force fields, including TIP3P-FB (52) water molecules. Solvent padding was 1 nm. Na^+^ or Cl^-^ ions were added to neutralize the system and create a 0.15 M ionic strength. Nonbonded forces were handled with the Particle Mesh Ewald (42) method, using a cutoff of 1 nm. The system was energy minimized with a 10 kJ/mol/nm minimization tolerance. Then, the system was equilibrated keeping a constant number of particles, volume and temperature (NVT system) in three steps, at 50 K, 150 K and 300 K with a 0.5 fs timestep, for 6000 steps at each temperature using the Langevin-Middle Integrator. Production steps were done in keeping a constant number of particles, pressure and temperature (NPT system), at 300 K and 2 fs timesteps. Bonds involving hydrogens were kept at a constant length. A Monte Carlo Barostat was used every 25 steps to keep a constant pressure of 1 bar and temperature of 300 K. For each system, a 100 ns long MD simulation was run. As each system contained four antibody-NA antigen pairs, a total of 400 ns was available to analyze for each antibody-NA antigen pair. The MDAnalysis (53, 54)python library was used to analyze the trajectories.

For testing the prediction capabilities of different AI-based structure prediction methods, we used the Boltz-2 (55), Chai-1 (18) and ESMFold2 (56). The VASP tetramerization domain – NA head sequences were given as input, in four copies to create a tetramer, along with one (Chai-1, ESMFold2, Boltz-2 with restraints) or four copies (Boltz-2 without restraints) of the Fv domains resolved for 2H08 and 3H03 in PDBs 8E6K and 8E6J respectively. Restraints were given as W329 – S32, D255 – Y55 and N387 – S106 between the VASP-N1 structures and the heavy chain Fv domain of 3H03 for Chai-1, and as W329 D255 and N387 of the VASP-N1 structures with the heavy chain Fv of 3H03. For 2H08, restraints were given as V65 – S104, N356 – T32 and N349 – Y54 between the VASP-N1 structures and the heavy chain Fv domain of 2H08 for Chai-1, and as V65, N356 and N349 of the VASP-N1 structures with the heavy chain Fv of 2H08. DockQ (57) was used to score the models, template was the input of the respective MD simulations, without including Ca^2+^-ions and glycosylations.

All models, MD simulation input files as well as final structures and scripts used for analyzing trajectories are available to download at ZENODO.

## Conflict of interest statement

The Icahn School of Medicine at Mount Sinai has filed patent applications relating to SARS-CoV-2 serological assays, NDV-based SARS-CoV-2 vaccines influenza virus vaccines and influenza virus therapeutics which list FK as co-inventor and FK has received royalty payments from some of these patents. Mount Sinai has spun out a company, Castlevax, to develop SARS-CoV-2 vaccines. FK is co-founder and scientific advisory board member of Castlevax. FK has consulted for Merck, GSK, Sanofi, Gritstone, Avimex, Curevac, Seqirus and Pfizer and is currently consulting for 3rd Rock Ventures. The Krammer laboratory is also collaborating with Dynavax on influenza vaccine development. VS is listed on the patent for the SARS-CoV-2 serological assay. The Simon and Krammer laboratories collaborate with Sanofi on SARS-CoV-2 vaccine strain selection.

## Data availability statement

All data underlying this manuscript have been deposited at Zenodo

## Acknowledgements

We thank Parul Singh and Joshua Yueh for help with protein expression and all the study participants for their generous support of our research programs. We also thank Lena Hansen and Ali Ellebedy for the initial isolation of the mAbs and Harm van Bakel and the Mount Sinai Pathogen Surveillance Program for providing virus isolates and sequences. Work at the Icahn School of Medicine at Mount Sinai was supported by institutional funds to C-VaRPP. Work at the Medical University of Vienna was supported by institutional funds to the Ignaz Semmelweis Institute.

